# Cell type specific polyploidization in the royal fat body of termite queens

**DOI:** 10.1101/2023.05.11.540364

**Authors:** Tomonari Nozaki, Eisuke Tasaki, Kenji Matsuura

## Abstract

Tissue-specific endopolyploidy is widespread among plants and animals and its role in organ development and function has long been investigated. In insects, the fat body cells of sexually mature females produce substantial amounts of egg yolk precursor proteins (vitellogenins) and exhibit high polyploid levels, which is considered crucial for boosting egg production. Termites are social insects with a reproductive division of labor, and the fat bodies of mature termite queens exhibit higher ploidy levels than those of other females. The fat bodies of mature termite queens are known to be histologically and cytologically specialized in protein synthesis. However, the relationship between such modifications and polyploidization remains unknown. In this study, we investigated the relationship among cell type, queen maturation, and ploidy levels in the fat body of the termite *Reticulitermes speratus*. We first confirmed that the termite fat body consists of two types of cells, that is, adipocytes, metabolically active cells, and urocytes, urate- storing cells. Our ploidy analysis using flow cytometry has shown that the fat bodies of actively reproducing queens had more polyploid cells than those of newly emerged and pre-reproductive queens, regardless of the queen phenotype (adult or neotenic type). Using image-based analysis, we found that adipocytes became polyploid during queen differentiation and subsequent sexual maturation. These results suggest that polyploidization in the termite queen fat body is associated with sexual maturation and is regulated in a cell type-specific manner. Our study findings have provided novel insights into the development of insect fat bodies and provide a basis for future studies to understand the functional importance of polyploidy in the fat bodies of termite queens.

## Background

Polyploid cells are widespread among various tissue and organs that exhibit high metabolic activity in plants and animals. They play pivotal roles in the regulation of gene expression, cell size, and differentiation [1–4]. Polyploid cells are generated by endocycle, or DNA amplification during the S phase of the cell cycle without subsequent cell division [2, 5]. Endocycles have been considered a low-cost strategy to increase cell and/or tissue size and to efficiently produce massive amounts of molecules needed during development, reproduction, immunity, and other life activities. This is because this process can avoid spending more time, materials, and energy to undergo normal mitosis [2, 5, 6]. However, there is still insufficient data connecting ploidy levels with cell specialization, gene expression levels/patterns, and organ development, which are required to determine the adaptive functions of tissue-specific polyploidy [6, 7].

The insect fat body is a multifunctional organ that is involved in the synthesis, storage, and secretion of lipids, proteins, and carbohydrates [8]. During oogenesis, vitellogenins or yolk protein precursors are massively produced in the female fat body and accumulate in developing oocytes [8, 9]. Sexually mature females of some solitary insects have more polyploid cells in their fat bodies than in other tissues or organs, which are most likely to boost vitellogenin production [10–12]. In social insects, which are characterized by a reproductive division of labor (Fig. 1A) [13], an intriguing relationship between tissue-specific polyploidy and the division of labor has been identified [14–18]. Nozaki and Matsuura [17, 18] demonstrated that in termites, high endopolyploidy only occurs in the queen fat body, but not in males or non-reproductive females. They also reported that highly fecund queens in foraging termite species had higher polyploid levels in their fat bodies than queens in wood-dwelling species, which were less fecund. These studies hypothesized that fat body polyploidy enhances egg production and is linked to histological and/or cytological modifications during reproductive maturation [19–23].

**Figure 1.**
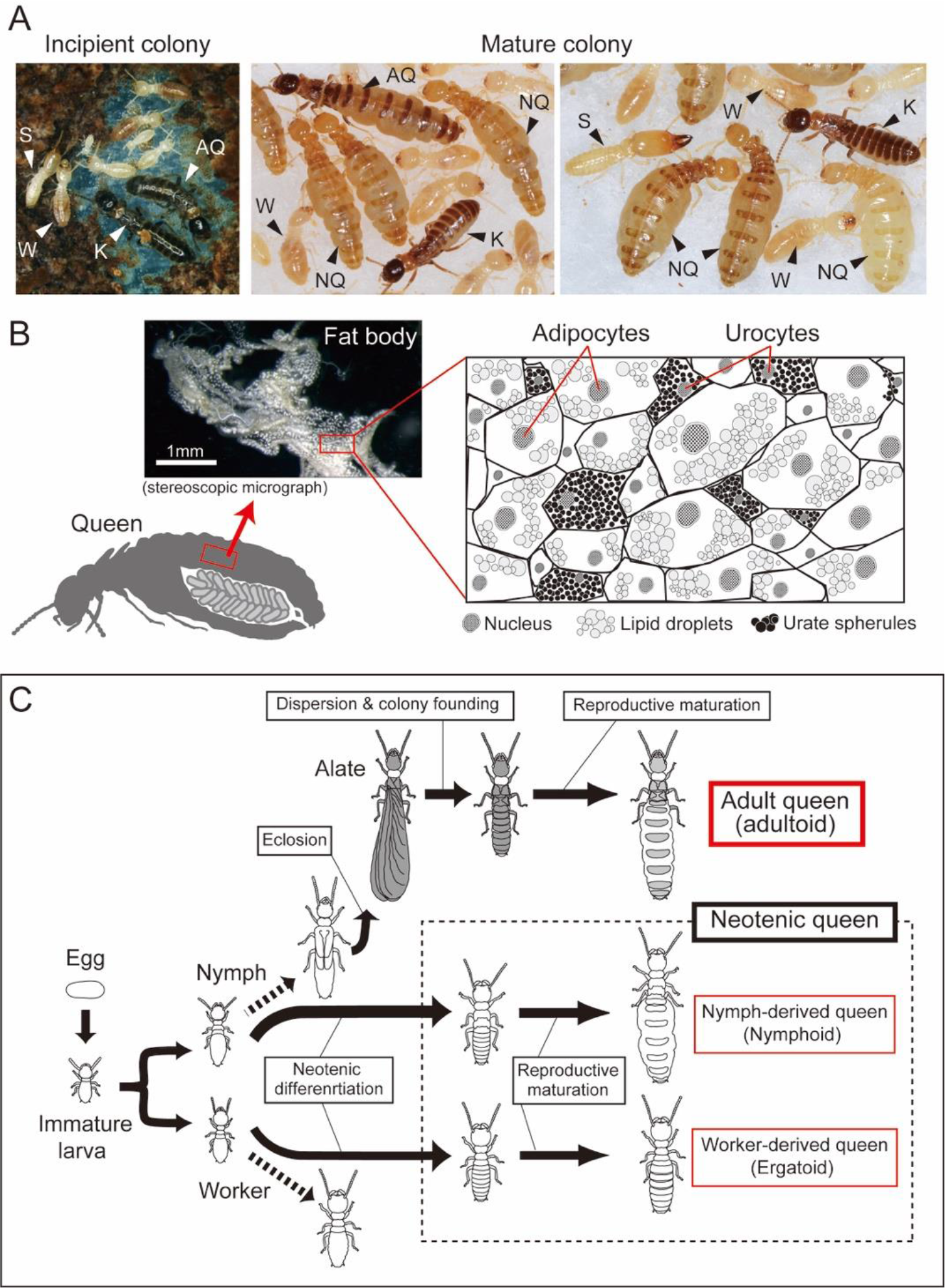
Queen development and the fat body structure in termites. **A** Photographs of *Reticulitermes speratus* colonies. Monogamous pairs of adults found new colonies. Mature field colonies comprise a number of queens, which continuously produce eggs over several years. AQ, adult queen; K, king; NQ, neotenic queen; S, soldier; W, worker. **B** The abdomen of a fully matured termite queen is almost filled with ovaries and fat body. In *R. speratus*, the fat body appears as whitish loose tissue located around the digestive tube and reproductive organs. The fat body tissue comprised two types of cells, that is, adipocytes, which are metabolically active and contain lipid droplets, and urocytes, which are not active but contain urate spherules. The illustration of the histology of the termite fat body was drawn with reference to [23]. **C** Developmental patterns of queens in *R. speratus*. Queens are categorized into adultoids (adult queens), nymphoids (nymph- derived neotenics) and ergatoids (worker-derived neotenics). All queens experience reproductive maturation after adult or neotenic molts. In the species, almost all queens collected in field colonies are nymphoid queens. Ergatoid queens are rarely observed in field colonies [24], yet they can be easily induced under laboratory conditions [25].

The termite fat body is composed of two types of cells, namely urocytes and adipocytes (Fig. 1B) [8, 23]. Urocytes are packed with urate spherules, which deposit uric acid as a nitrogen reservoir. Adipocytes are metabolically active cells that synthesize and store lipids, glycogen, and various proteins including vitellogenins. Only the primitive termite *Mastotermes darwiniensis* has a third cell type, that is, bacteriocytes, that harbors the bacterial symbiont, which is ubiquitous in cockroaches [23, 26]. The fat bodies of functional queens contain more adipocytes than urocytes, and the adipocytes of fully mature queens specialize in the synthesis of proteins [23]. In higher termites, the fat body of queens contains abundant proteins and RNAs, which has shown considerable differences from that of nonreproductives (royal fat body; [19, 20]). Based on these observations, adipocytes are likely to show higher ploidy levels than urocytes. However, to date, there has been no information on the cellular specificity of fat body polyploidization. Cytological information on ploidy dynamics could provide further implications for the functional significance of fat-body polyploidy.

*Reticulitermes* is a well-studied termite genus in terms of its reproductive biology, oogenesis or ovarian development, and caste differentiation (Fig. 1C) [27–31], and the complete genome has recently been provided [32]. In *R. speratus*, most mature field colonies contain tens to hundreds of neotenic queens [33, 34], which are usually derived from nymphs [24]. Meanwhile, nymph-derived and worker-derived neotenics can be induced under laboratory conditions [35, 36]. Adult queens (derived from imago) are occasionally collected from field colonies [24], and colony initiation using collected alates under laboratory conditions is frequently conducted by many researchers (Fig. 1A) [30, 37, 38]. Therefore, this species is suitable for comparative studies of queen types and their maturation levels.

In this study, we investigated the relationship between cell type and ploidy level in the fat body of the termite *R. speratus*, with a particular focus on queen differentiation, reproductive maturation, and fecundity. We first conducted microscopic observations of the cytological features of the termite fat body and confirmed the presence of two types of cells, that is, adipocytes, which contain many lipid droplets, and urocytes, which have many spherical crystals surrounding the nucleus. It was also confirmed that vitellogenin genes were highly expressed in the fat body of queens using quantitative RT-PCR. Using flow cytometry, we examined the ploidy dynamics in the fat bodies of three types of termite queens in the species, that is, nymph- or worker-derived neotenics and adult queens. Our image-based analysis of nuclear size and ploidy levels showed cell specificity in the polyploidization of termite queen fat bodies. Based on our results, we have discussed the functional and developmental importance of fat body polyploidization during queen maturation and fecundity.

## Methods

### Microscopic observation of adipocytes and urocytes in the termite fat body

To visualize the differences between adipocytes containing lipid droplets and urocytes with accumulation of urate crystals, we performed a morphological analysis of the fresh fat body of *R. speratus* using a confocal laser-scanning microscope (FV1000, Olympus). The fat bodies of workers and late-instar nymphs, collected from pine or cedar forests in Aichi, April 2021, were dissected in phosphate-buffered saline (PBS; 33 mM 143 KH2PO4, 33 mM Na2HPO4, pH 6.8). This process was conducted with fine forceps under a stereomicroscope (Olympus SZX7, Olympus). The fat body of the termites appeared as whitish loose tissue located around the digestive tube and reproductive organs (Fig. 1B) [23]. We dissected the fat body from the abdomen of the insects with care to avoid contamination by other tissues, such as Malpighian tubules and tracheoles. The dissected fat bodies were treated with trypsin buffer (0.25% trypsin [Nacalai Tesque] in PBS) for 15 min to loosen the adhesion between the cells. The tissues were stained using Hoechst 33342 (1 μg/mL, Dojindo) and Lipi-Red (1 nmol/mL, Dojindo) in PBS for 15 min. The tissues were directly mounted on a glass slide, crushed using a cover slip and visualized using fluorescent and differential interference contrast (DIC) microscopy. Note that we first tried to observe fat bodies fixed by 4% PFA, permeabilized by PBS-T, and stained with DAPI. However, this was not suitable because no urate crystals were observed, potentially because of the dissolution of urates during sample processing (data not shown). We also conducted histological observations of the fat bodies of workers and queens collected in August 2018 from a field colony in Kyoto in August. The dissected fat bodies were fixed overnight in PBS with 2% glutaraldehyde and 4% paraformaldehyde at 4 °C and postfixed with 1% OsO4 for 2 h. The samples were then dehydrated and embedded in epoxy resin. Ultrathin sections were stained with 2% saturated uranyl acetate solution and 2.5% lead citrate solution and examined using transmission electron microscopy (Hitachi-7650, Hitachi). The electron microscopy study was supported by the Division of Electron Microscopy, Center for Anatomical Studies, Graduate School of Medicine, Kyoto University.

### Ploidy analysis of the fat body of termites using flow cytometry

To investigate the ploidy dynamics in the fat bodies of queens in *R. speratus*, we compared the ploidy levels of fat bodies among queens at different maturation stages. This analysis was conducted for all three queen types in the species, that is, nymph- derived neotenic (nymphoids), worker-derived neotenic (ergatoids), and alate (adultoid) queens (Fig. 1C). Details of the three types of termite queen are presented in the Supplementary Information (SI Methods and Table S1). The fresh body weights of individual termites were measured using a digital balance and expressed in milligrams to two decimals. Fat bodies were dissected in PBS using fine forceps under a stereomicroscope (Olympus SZX7; Olympus). We used the heads of all the individuals analyzed as diploid tissues [17, 18]. Tissues were processed for flow cytometric analysis using the Cycletest PLUS DNA Reagent Kit (Becton Dickinson). All the procedures were performed according to [17, 18]. Nuclei were stained using propidium iodide (PI) and analyzed for DNA-PI fluorescence with an Accuri C6 Flow Cytometer (Becton Dickinson) at an excitation wavelength of 488 nm. Approximately 1,000 cells were acquired for each measurement. Gating was performed using Accuri C6 software v1.0.264.21 to measure the number of nuclei at each ploidy level (2C, 4C, and 8C) per sample. Representative histograms are shown in Fig. 2.

### Analysis of ploidy, cell types, and the size of nuclei in the fat body using an image-based method

To infer the cell type specificity of polyploidization in termite fat bodies, we first confirmed the relationship between the ploidy level and size of nuclei using a cell sorting technique and microscopic observation. Nuclei sorted from the head and fat body of queens, according to each ploidy level (2C, 4C, and 8C), were measured (see SI Methods). Using semi-destructive treatment and microscopy, we compared the nuclear size, as a proxy for ploidy, between two types of cells, adipocytes and urocytes, in the fat bodies of workers, nymphs, and young and mature nymphoid queens. The fat bodies of workers and nymphs from two field-collected colonies (HM210907A and OZ210902) and young and reproducing nymphoid queens from lab-orphaned colonies derived from the colonies were dissected in PBS (for details see Table S1). For each colony, one individual of each termite type was randomly selected (n = 2 for each type). The tissue was treated with trypsin buffer (0.25% trypsin in PBS) for 10 min to loosen the adhesion between cells. The tissues were stained with Hoechst 33342 in PBS (1 μg/mL, Dojindo) for 10 min. The stained tissues were directly mounted on a glass slide, crushed using a cover slip, and visualized using fluorescent and differential interference contrast (DIC) microscopy. Nuclei were randomly captured using a DS-Fi1 CCD camera (Nikon, Japan). The cell type and their number were recorded based on the DIC images. The size of the nuclei was measured using Hoechst fluorescent images, with the image analysis software ImageJ (National Institutes of Health, USA, http://rsb.info.nih.gov/ij/).

### Statistical analysis

To examine ploidy dynamics in queen differentiation and reproductive maturation, the proportion of polyploid cells in the fat body, which was calculated by the nuclei count with value of 2C as diploid, and 4C, 8C, and higher as polyploid, were compared among different maturation stages of nymphoid, ergatoid, and adultoid queens, and the ones from which they were derived, namely nymphs, workers, and alates. We used the worker data presented in [18]. The proportion of polyploid cells in the fat body of each individual was analyzed using generalized linear mixed-effect models (GLMM) with binomial errors and a logit-link function. This was then followed by the Tukey’s HSD post hoc test. In this analysis, the category of individuals, that is, nymphs, nymphoid 0 days, and reproducing nymphoid queens, and body weight were treated as fixed effects, and the original colony was included as a random effect. To infer the relationship between the fecundity and ploidy levels of their fat bodies, that is, to determine whether the endopolyploidy level could be positively correlated with the egg production capacity of queens, we conducted a correlation analysis of the proportion of polyploid cells in the fat body and the fresh body weight of queens. This was used as a proxy of fecundity (Results and SI methods) using GLMM with binomial errors and a logit-link function. For this analysis, we used data from mature nymphoid queens collected from field colonies. In this model, the proportion of polyploid cells in the fat body of queens was the response variable, and the initial fresh body weight was the explanatory variable. The original colony was included as a random effect. To examine whether the polyploidization pattern was different between the cell types (adipocyte/urocyte) in the fat body of termites, and to determine the relationship between cell type and ploidy levels, the size of nuclei, that is, a proxy of ploidy level, (Results and SI methods) from workers, nymphs, young, and reproducing nymphoid queens were compared between adipocytes and urocytes. In this analysis, we used LMM followed by Tukey’s HSD post-hoc test, wherein the cell type and category of individuals were fixed effects, and the original colony was a random effect. All the analyses were conducted using the “car,” “emmeans,” “lawstat,” “lme4,” and “multcomp” packages in R software v4.1.1 (https://www.r-project.org/). Additionally, “beeswarm” and “vioplot” were used to draw the beeswarm and violin plots, respectively. Differences were considered significant when the *p* value was < 0.05.

## Results

### Morphological observation of the termite fat body

Consistent with previous observations in other termites [19, 20, 23], the fat body of *R. speratus* consisted of cells that contained numerous lipid droplets and urate spherules, corresponding to adipocytes and urocytes, respectively (Fig. S1). Confocal microscopy showed that the nuclei of adipocytes were circular and varied in size, whereas those of urocytes were slightly deformed and small (Fig. S1A). The TEM observations were consistent with those of previous reports on other termite species. Two types of cells were recognizable, that is, cells with lipid droplets or urate crystals (Fig. S1B). Based on these observations, we cytologically confirmed the presence of termite adipocytes and urocytes (Fig. S1C).

### Vitellogenin gene expression in the fat body of termites

Three vitellogenin genes (*RsVg1*–*3*) were highly expressed in queen fat bodies (Fig. S2). Among the fat bodies of females, queens always exhibited significantly higher expression levels than the workers and soldiers (Mann–Whitney U test with Bonferroni adjustment, *p* < 0.05). In males, the expression levels of vitellogenin genes were generally relatively low. However, *RsVg2* was highly expressed in the fat body and testes of two of the five kings.

### Ploidy dynamics in the termite queen fat body

#### Nymphoid queens

Among the field collected nymphoid queens, we recognized “young” and “matured” queens, as shown in the Methods. While diploid cells were dominant in the fat bodies of young queens, tetraploids (i.e., 4C cells) were always the most abundant ploidy class in the fat bodies of mature queens (Fig. 2A). The fat body of mature nymphoid queens contained significantly more polyploid (2C, 4C, 8C and 16C) cells than those of the young queens (GLMM with type II Wald chi-square test, types of queens: *χ*^2^ = 1280.49, *df* = 1, *p* < 0.01, Fig. 2B). There were also significant effects of fresh body weight on the proportion of polyploid cells in fat bodies (body weight: *χ*^2^ = 154.93, *df* = 1, *p* < 0.01). Among the female nymphs and laboratory-induced nymphoid queens, there were significant differences in the proportion of polyploid cells in the fat body (GLMM with type II Wald chi-square test, female category: *χ*^2^ = 139.0634, *df* = 2, *p* < 0.01; Fig. 3). Body weight did not significantly affect the degree of polyploidy (body weight: *χ*^2^ = 0.0101; *df* = 1; *p* = 0.92). Reproducing nymphoids exhibited the highest proportion of polyploid cells in their fat bodies, and newly molted nymphoids showed significantly lower values than nymphs (Tukey’s HSD, *p* < 0.01, Fig. 3). In all individuals, the head cells predominantly consisted of diploid cells.

**Figure 3.**
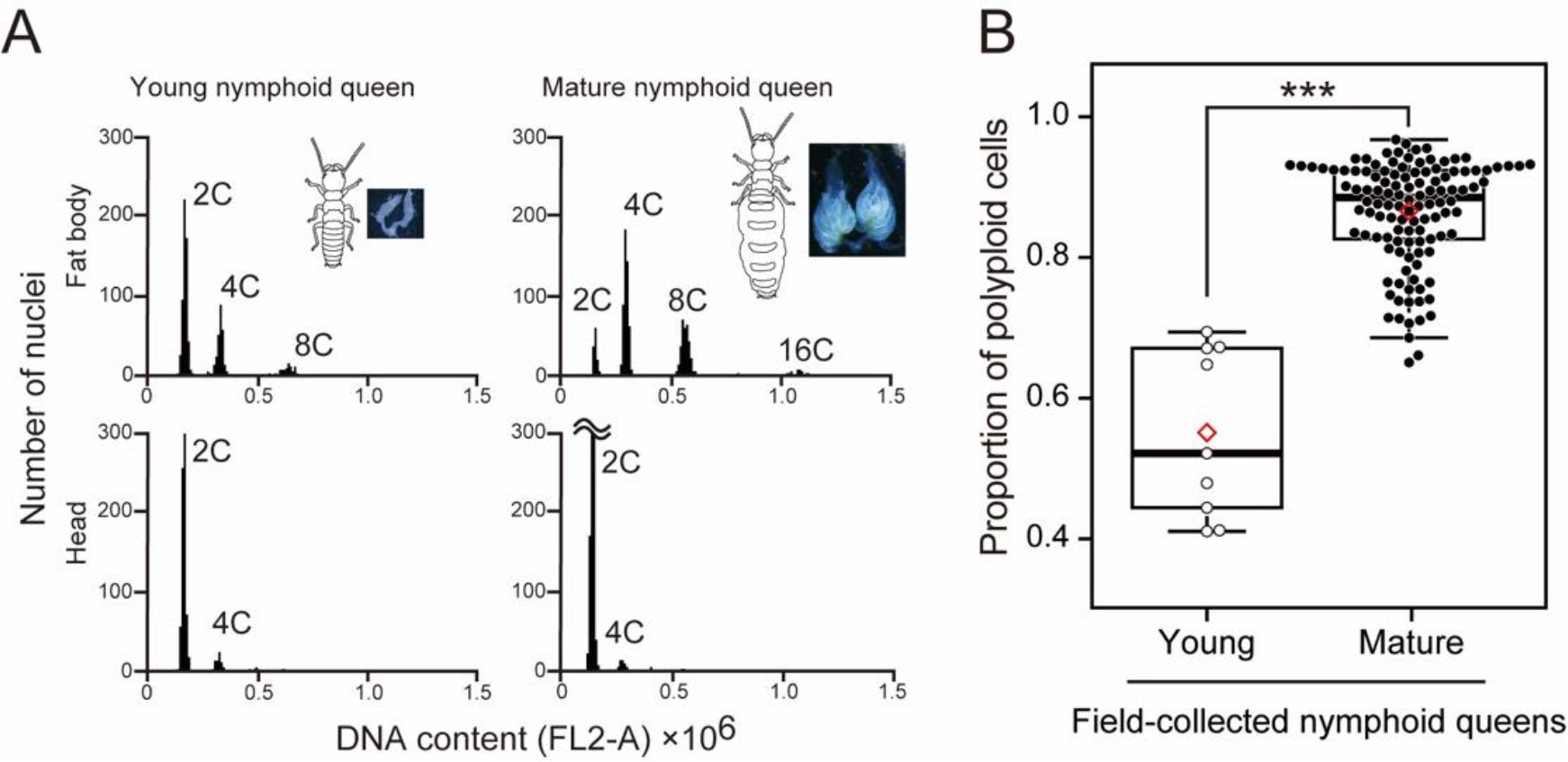
Ploidy analysis from flow cytometry of the fat body and head cells from young and matured queens from field colonies. **A** Examples of the result histograms from young and matured queens are presented. In each histogram, the first peak corresponds to the distribution of 2C-DNA nuclei, whereas the second, the third and the fourth correspond to the distribution of 4C- 8C- and 16C DNA nuclei, respectively. The ploidy was determined by analysis of the head cells (diploid; 2C, [17]). Images of insects and ovaries were included. **B** The proportion of polyploid cells in the fat body between young and mature field collected nymphoid queens. Box plots show the median (center line), 75th percentiles (top of box), 25th percentiles (bottom of box), and whiskers connect the largest and smallest values within 1.5 interquartile ranges. The red diamonds indicate mean percentages of each category. Dots are individual values. Asterisks indicate a significant difference (linear mixed effect model with a Wald chi-square test, \*\*\**p* < 0.05).

#### Ergatoid queens

Among the female workers, newly molted ergatoids, and reproducing ergatoid queens, there were significant differences in the proportion of polyploid cells in the fat body (GLMM with type II Wald chi-square test, female category: *χ*^2^ = 18.918, *df* = 2, *p* < 0.01; Fig. 3), considering the significant effect of body weight (body weight: *χ*^2^ = 35.265, *df* = 1, *p* < 0.01). Reproducing ergatoid queens had a higher proportion of polyploid cells in their fat bodies than newly molted ergatoids and female workers (Tukey’s HSD, *p* < 0.01, Fig. 3). In all individuals, the head cells predominantly consisted of diploid cells.

#### Adultoid queens

There were originally four categories for this type of queen, that is, female alate, field-collected founding queen, founding queen in laboratory-established colonies, and field-collected fully matured queens. However, we found that the results from field-collected founding queen and founding queen in laboratory-established colonies were similar. Therefore, we pooled them as “founding adultoids.” Among the female alates, founding adultoids, and fully mature queens, there were significant differences in the proportion of polyploid cells in the fat body (GLMM with type II Wald chi-square test, female category: *χ*^2^ = 226.84, *df* = 2, *p* < 0.01; Fig. 3), considering the significant effect of body weight (body weight: *χ*^2^ = 196.29, *df* = 1, *p* < 0.01). Fully matured adultoid queens had the highest proportion of polyploid cells in their fat bodies, with founding queens showing significantly lower values than the female alates (Tukey’s HSD, *p* < 0.01, Fig. 3). In all individuals, the head cells predominantly consisted of diploid cells.

### Polyploid levels in the fat body and fecundity of nymphoid queens

We examined the relationship between fresh body weight and egg production rate for five days, and found a significant effect of fresh body weight on the number of eggs oviposited by queens over five days (GLMM with type II Wald chi-square test, *χ*^2^ = 97.308, *df* = 1, *p* < 0.01, Fig. S4). Therefore, we used the fresh body weight of the queens as a proxy for fecundity. Focusing on the polyploid cells in the fat bodies of field-collected and reproducing nymphoid queens, we inferred a relationship between fecundity using fresh body weight, and the level of endopolyploidy. We found that fresh body weight had a significant effect on the proportion of polyploid cells in the fat bodies of queens (GLMM with type II Wald chi-square test, fresh body weight: *χ*^2^ = 109.71, *df* = 1, *p* < 0.01). However, the effect was negative, and the correlation value was relatively low (Fig. S4 and S5).

### Cell-type specific polyploidy in the fat body of termite queens

We assessed the relationship between nuclear size (area) and ploidy level. There was a significant difference in size among 2C nuclei sorted from the head sample and 2C, 4C, and 8C nuclei sorted from fat body samples (Fig. S6; GLMM with type II Wald chi-square test, nuclear category: *χ*^2^ = 431.94, *df* = 3, *p* < 0.01). Although there were some overlaps, nuclei with higher ploidy levels exhibited significantly larger sizes (Tukey’s HSD: 2C from head = 2C < 4C < 8C from fat body, *p* < 0.01; Fig. S6). This clear correlation allowed us to use nuclear size as a proxy for the ploidy level.

Based on DIC microscopy, we confirmed that we could morphologically distinguish between the two cell types. Nuclei surrounded by urate spherules and lipid droplets were categorized into urocytes and adipocytes, respectively (Fig. 4A). There were significant differences in the proportions of adipocytes and urocytes. Reproducing nymphoids had more adipocytes (89.8%) than workers (76.7%), nymphs (78.8%), or newly molted nymphoids (76.1%) (Fisher’s exact test with Bonferroni correction, *p* < 0.001). We then investigated the relationship between cell type and nuclear size and found that both cell type and individual type had significant effects on nuclear size (LMM with type II Wald chi-square test, cell type: *χ*^2^ = 156.357, *df* = 1, *p* < 0.01; individual type: *χ*^2^ = 501.615, *df* = 3, *p* < 0.01). This interaction was significant (*χ*^2^ = 58.853, *df* = 3, *p* < 0.01). The size of adipocytes significantly increased with queen differentiation and maturation (Tukey’s HSD; worker = nymph < newly molted nymphoid < reproducing nymphoid, *p* < 0.01, Fig. 4B). Meanwhile, there was no significant difference in the size of urocytes among workers, nymphs, and queens (Tukey’s HSD, *p* < 0.01, Fig. 4B). In both cell types, the size variation was relatively large, regardless of termite phenotype and developmental stage, as suggested in the analysis of the sorted nuclei (Fig. S6).

**Figure 4.**
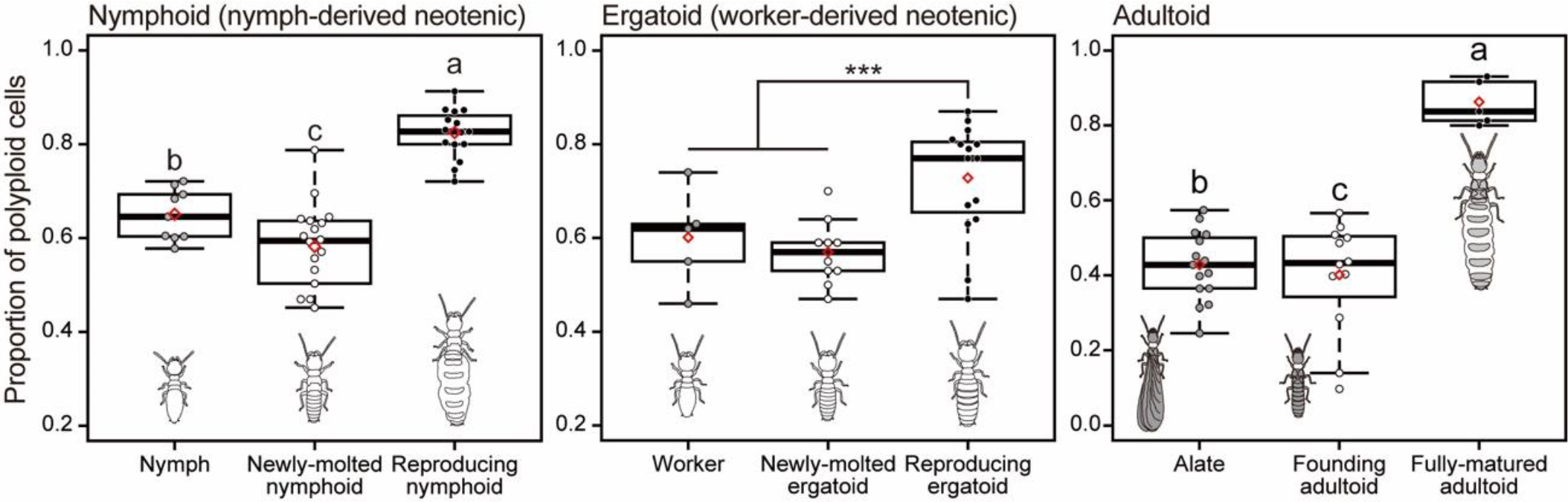
The proportion of polyploid cells in the fat body along with development of three types of queens. Box plots show the median (center line), 75th percentiles (top of box), 25th percentiles (bottom of box), and whiskers connect the largest and smallest values within 1.5 interquartile ranges. The red diamonds indicate mean percentages of each category. Dots are individual values. Different letters and asterisks indicate significant differences among means in each comparison group (Tukey’s HSD test, *p* < 0.05).

**Figure 5.**
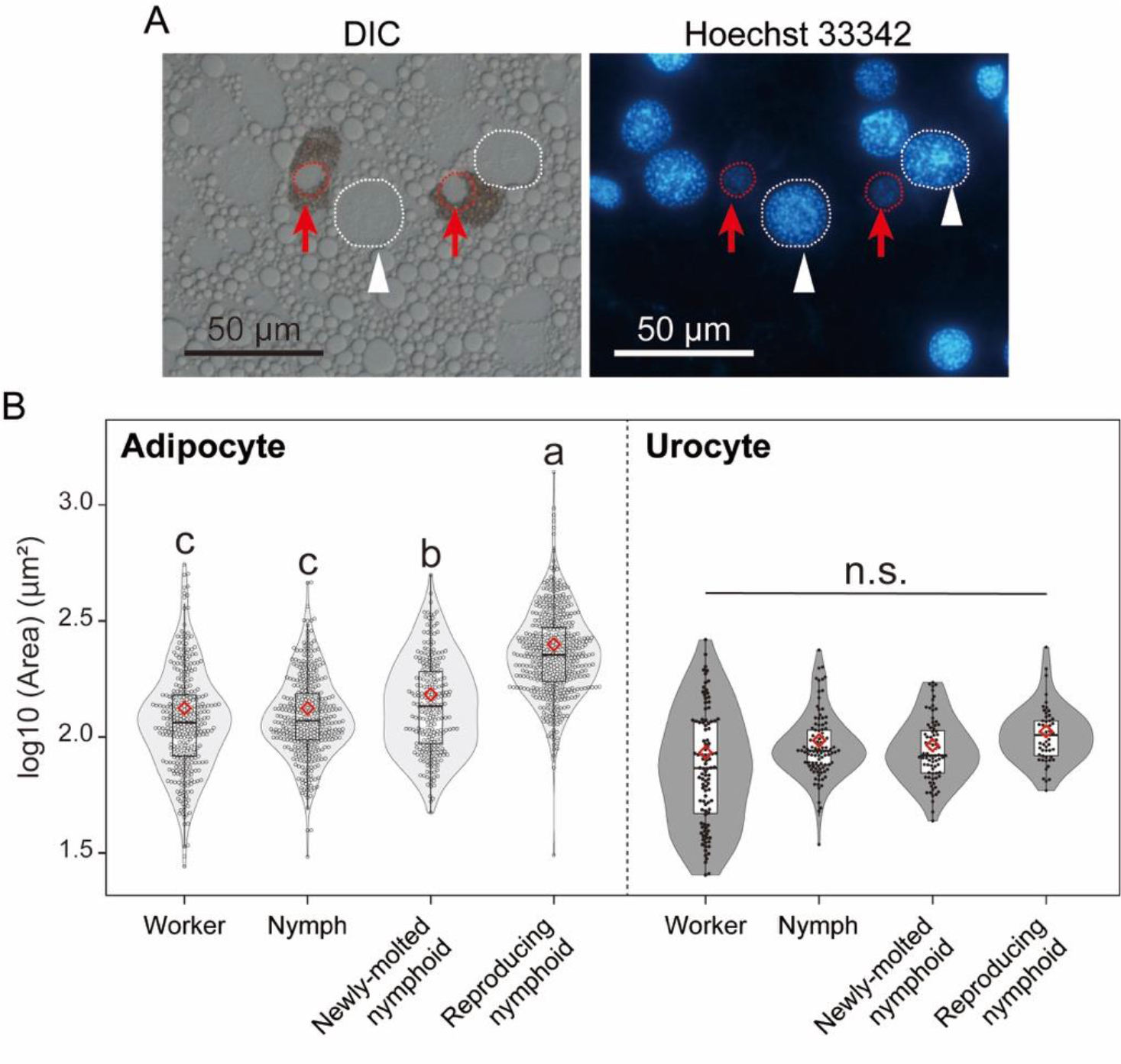
The relationship between cell types and nuclear size (a proxy of ploidy level) in the fat body of *R. speratus*. **A** Under DIC, adipocytes can be discriminated from urocytes, because adipocytes were not surrounded by urate spherules. Nuclei of both types of cells were stained and visualized by Hoechst 33342. White arrowheads indicate representative nuclei of adipocytes (enclosed by white dashed lines). Red arrows indicate urocyte nuclei (enclosed by red dashed lines). **B** Size distribution of nuclei of adipocytes and urocytes in the fat body of worker, nymph and immature and mature nymphoid queens. Violin plots showing the distribution of the size (area) of nuclei. Box plots show the median (center line), 75th percentiles (top of box), 25th percentiles (bottom of box), and whiskers connect the largest and smallest values within 1.5 interquartile ranges. The red diamonds indicate mean size of each category. Dots are individual values. Different letters indicate significant differences among means in each comparison group (Tukey’s HSD test, *p* < 0.05).

## Discussion

In the present study, we have demonstrated that during the maturation of termite queens, their fat bodies become polyploid in a cell type-specific manner. Our flow cytometric analysis of field-collected (Fig. 2B) and laboratory-grown queens (Fig. 3) showed that the ploidy levels of the fat bodies were higher in reproducing queens that had fully developed ovaries than in young or newly molted ones, as recently shown in a higher termite [39]. This pattern was maintained regardless of the queen phenotype, that is, nymph-derived neotenic, worker-derived neotenic, and adult queens (Fig. 3). We also found that adipocytes increased their ploidy levels in the fat bodies of termite queens (Fig. 4B). In adipocytes, nuclear size, which is a proxy for ploidy levels (Fig. S5) increased significantly during queen maturation. In contrast, the size of urocyte nuclei was not significantly different between female workers and young/mature queens (Fig. 4B). The fat body highly expressed vitellogenin genes (Fig. S2), and “adipocyte” was assumed to be a principal cell type in metabolic activity of the fat body [8, 23]. Therefore, these results were compatible with the idea that the polyploidization in the fat body could contribute to increasing egg production or massive vitellogenin synthesis [18]. In future studies, gene expression, cell types, and ploidy levels should be examined in detail, alongside cell sorting techniques and single-cell-level gene expression analysis.

The dynamics of the ploidy level and the proportion of cell types that we described in the present study are consistent with previous histological studies on the development of “royal fat body [19, 20].” The fat bodies of functional queens in higher termites contain a larger number of adipocytes than urocytes, and the adipocytes of fully matured queens are specialized in the synthesis of proteins. They have limited lipid content but numerous rough endoplasmic reticulum and Golgi apparatus [23]. Specialization of adipocytes through polyploidization may be essential for the formation of “royal fat body” in termite queens. The relationship between royal fat body development and polyploidy should be addressed using integrated approaches, including cytological, histological, and omics analyses.

There have been several reports on age-related polyploidy in insect tissues and organs [14, 40]. However, this study has shown that increasing the ploidy level in the fat body of termite queens is closely linked to reproductive maturation rather than being age-related. This is because three months after emergence from artificially established colonies, queens exhibited almost the same degree of ploidy in their fat bodies as field-collected queens that would otherwise have been bred for several years (unknown exact age) (Fig. 2B, Fig. 3). Data from a higher termite also suggested that ploidy levels in the fat bodies of queens increased sharply during the early stages of reproductive maturation [39]. Therefore, rather than aging promoting fat body polyploidy per se (the possibility cannot be ruled out), fat body polyploidy is likely to play an essential role in high reproductive outputs and the extraordinarily long lives of termite queens, such as increasing egg production or prevention of aging [41, 42].

Juvenile hormone (JH) promotes fat body cell polyploidization in locusts [10, 11, 43–45]. In termites, JH plays critical roles in neotenic and soldier differentiation [46–48]. In general, insect JH is important for ovarian maturation and oogenesis [49]. Therefore, polyploidy in the fat body of neotenic queens is likely attributed to neotenic differentiation with increasing JH titers. Nevertheless, we found that polyploidization of the termite fat body occurred during queen maturation rather than during neotenic differentiation (Fig. 3). The fat bodies of newly molted neotenics [nymphoids (58%) and ergatoids (57%) had a similar proportion of polyploid cells as nymphs (65%) and workers (60%). However, these proportions were significantly different from those of reproducing neotenics including nymphoids (82%) and ergatoids (73%) (Fig. 3). Alates (42%) and founding (young) adult queens (40%) had a lower proportion of polyploid cells in their fat bodies than fully mature adults (86%) (Fig. 3). After initiation of the colony, female alates (founding queens) oviposit a small number of eggs for a month [38, 50]. This indicated that high-level polyploidy in the fat body is not essential for oogenesis. Fat body polyploidization may be critical for the mass production of eggs, although this should be assessed more directly in future studies.

In conclusion, our results have shown that the patterns of polyploidization and cell-type specificity in the fat body of termite queens, which is likely linked to queen fecundity [17, 18, 39]. Our data has also emphasized that fat body polyploidy was associated with queen maturation, that is, “the formation of royal fat body,” rather than being from aging or caste differentiation. Therefore, it is necessary to investigate whether endopolyploidy plays a role in increasing or regulating the transcription of rRNA and mRNA per nucleus, as shown in plants [7]. Our study has provided a foundation for further molecular and single-cell level analyses of the functional roles and proximate mechanisms of polyploidy in insect fat bodies.

## Declarations

### Availability of data and materials

The raw data and materials are available upon request from the authors.

### Ethics approval and consent to participate

Not applicable.

### Consent for publication

Not applicable.

## Author contributions

T. N. and K. M. designed the research; T. N. and E. T. performed the experiments and analyzed the data; T. N., E. T., and K. M. wrote the original draft of the paper; and all authors contributed substantially to revisions.

## Acknowledgements

We thank M. Takata, N. Mizumoto, T. Inagaki, T. Fujita, N. Yoshioka, and other members of the laboratory of Insect Ecology at Kyoto University for sample collection. We appreciate N. Hirose for experimental assistance, S. Shigenobu for technical advice, the Functional Genomics Facility, and the NIBB Core Research Facilities for technical support. We would also thank to Editage (www.editage.com) for English language editing.

