## Additional file 1 for "Cell type specific polyploidization in the royal fat body of termite queens"

Original Research Article

Supplementary Information

Tomonari NOZAKI^1, 2^, Eisuke TASAKI^1, 3^, Kenji MATSUURA^1^

^1^*Laboratory of Insect Ecology, Graduate School of Agriculture, Kyoto University, Kyoto, Japan*

^2^*Laboratory of Evolutionary Genomics, National Institute for Basic Biology, Okazaki, Japan*

^3^*Department of Biology, Faculty of Science, Niigata University, 8050 Ikarashi 2-no-cho, Nishi-ku, Niigata 950-2181, Japan*

*Correspondence:* Tomonari Nozaki, Laboratory of Evolutionary Genomic, National Institute for Basic Biology, Okazaki 444‐8585, Japan

Email:

 (TN)

 (ET)

 (KM)

ORCID :

https://orcid.org/ 0000-0003-4358-8118 (TN)

https://orcid.org/0000-0002-6514-5134 (ET)

https://orcid.org/0000-0002-9099-6694 (KM)

**SI Methods**

**Vitellogenin gene expression analysis by quantitative PCR**

To quantitatively determine vitellogenin synthesis in the queen fat body of *R. speratus*, the expression levels of three vitellogenin genes (*RsVg1*–*3*) [51] were compared between queen body parts, that is, the fat body and ovaries. The fat bodies of other colony members, namely the workers, soldiers, and kings, and the testes of kings were included in this experiment as negative controls for vitellogenin gene expression.

Five *R. speratus* colonies containing queens and kings were collected from secondary forests in Yamaguchi and Shiga, Japan, from May to June of 2016 and 2017. Rotten wood containing their colonies was transported to the laboratory and carefully dissected to extract the termites. The ovaries of the queens, testes of the kings, and fat bodies of all members were dissected in PBS with fine forceps under a stereomicroscope (Olympus SZX7, Olympus). Isolated tissues were temporarily preserved at −80 °C. Females and males were separated based on the shape of the caudal sternite using a stereoscope [52, 53]. We used nymph-derived (nymphoid) queens and adultoid kings as “queens” and “kings” in this study, respectively. This is because of the rare appearance of adultoid queens and neotenic kings of this species in nature [33]. Using an RNeasy mini kit (Qiagen), the total RNA was extracted individually from the fat bodies of workers, soldiers, and reproductive workers, including neotenic queens and adult kings. For the workers and soldiers, three individuals were pooled per sample. The ovaries of queens and the testes of kings were also included. cDNA was synthesized from the extracted RNA, using a PrimeScript™ RT reagent kit (Takara) and preserved at −20 °C. qPCR was performed using an Applied Biosystems® StepOne™ system (Thermo) with Power SYBR™ Green PCR master mix (Thermo). All the procedures were performed according to the manufacturer’s instructions. Detailed information on the vitellogenin genes (*RsVg1*–*3*) and primers for qPCR has been described previously [51]. Nicotinamide adenine dinucleotide dehydrogenase subunit 5 (ND5) was selected as the reference gene. Relative expression levels were calculated using a typical ΔΔCt method relative to the fat body of female workers. In total, we performed five replicates for each comparison group, including the queen ovaries, king testis, and fat body of the queen, king, (female and male) workers, and (female and male) soldiers. Each replicate was derived from a single colony, with five colonies in total. For the statistical analyses of the qPCR experiments, we used the Mann–Whitney U test with Bonferroni correction for multiple comparisons. We compared the relative expression levels in the fat bodies of female termites, that is, the queens, workers, and soldiers.

**Three types of termite queens for ploidy analysis by flow cytometry**

In total, 36 field colonies of *R. speratus* were collected from pine and cedar forests in Japan from April to September in 2015, 2016, and 2018. Of these, 21 colonies contained nymphoid or adultoid queens. Eleven colonies that did not contain royals, that is, satellite colonies, were used to establish artificial colonies and to obtain newly differentiated neotenics. The remaining three colonies without royals contained alates that were used as colony foundations to obtain young adultoid queens (see paragraphs below). Among the field-collected queens, most were nymphoids and a few were adultoids (Fig. 1A). As previously reported, no ergatoid queens were collected [31, 32]. Detailed information on the colonies used in the analyses is provided in Table S1.

For *R. speratus*, we obtained 127 nymph‐derived neotenics (nymphoids). They were categorized into “mature” and “young” field collected nymphoid queens based on their ovarian status. Mature queens had vitellogenic or pre-vitellogenic oocytes and, to a greater or lesser extent, a prominent yellow body at the base of the ovarioles, which is evidence of oogenesis [52]. The young queens did not have any vitellogenic or pre-vitellogenic oocytes in their ovaries or yellow bodies. In *R. speratus*, queens are cyclically replaced through parthenogenesis. Therefore, colonies of young queens could be collected [32]. In the present study, two of the 14 colonies contained young and mature queens, 13 colonies contained only mature queens, and one colony contained only young queens (Table S1). In total, we used 118 mature and nine young nymphoid queens for ploidy analysis of the fat body.

In the absence of reproductives, *Reticulitermes* workers and nymphs can develop into juvenile “neotenic” reproductives (Fig. 1C) [50]. To obtain laboratory-induced nymphoid and ergatoid queens, we established two types of orphan colonies using termite colonies without royals, that is, small-scale orphan colonies, wherein neotenic differentiation can be checked daily, and large-scale orphan colonies, in which termites can be provided with enough food to actively produce eggs, while we cannot check the details of colony members (Fig. S3). To obtain the young stages of neotenic queens, 100 workers with/without 10 late-instar nymphs from five colonies (YS180815A, YS180815B, YO180916, YO151006, and UR151008) were placed in a 30-mm Petri dish lined with moist unwoven cloth (small-scale orphan colonies, Fig S3). The dishes were maintained at 25 °C in the dark. The colonies were checked daily, and female neotenics, that is, 0-day nymphoid and egratoid queens, were collected. They were dissected on the same day and used for ploidy analysis. In colonies without nymphs, workers can differentiate into ergatoid neotenic queens, whereas in colonies with 10 nymphs, only nymphoid queens are usually observed [31, 50]. When male neotenics emerged, they were removed and not used. Three late-instar nymphs from the colonies YS180815A, YS180815B, and YO180916 were included in the ploidy analysis. However, it was relatively difficult to maintain healthy neotenic queens in small-scale orphan colonies. Almost all the queens were attacked and killed by workers after one or three months. Therefore, we established large-scale orphan colonies, wherein approximately 3000 workers were with/without 100 nymphs from five colonies (HI150720, UR150715A, YO150715B, B150703, and OK150625B) (Fig. S3). The termites were placed in a plastic case (10 × 10 × 3 cm) filled with mixed sawdust bait and kept at 25 °C in constant darkness for three months. Immediately before the experiment, the baits were carefully dissected to extract the individuals. Egg piles were observed in all large-scale orphan colonies, indicating that the emerged nymphoid or ergatoid queens were actively reproducing. After three months, we used the queens for ploidy analysis of the fat body, with confirmation of the ovarian status. Detailed colony information used in the rearing induction of neotenics is provided in Table S1.

To obtain adultoid queens with different stages of reproductive maturation, we collected termite colonies containing alates, and established founding colonies according to [54]. Alates that shed their wings were chosen randomly from three colonies, and three female alates from each colony were used for ploidy analysis. Female–male pairs were established with individuals from the same colony, and each pair was placed in a 55 mm plastic Petri dish containing mixed sawdust bait blocks. The dishes were maintained at 25 °C in the dark. After approximately three months, the bait blocks were carefully destroyed, and the founding mating pairs, that is, the primary queens and kings, were extracted. We also collected three young founding colonies from a field containing female and male dealates and eggs. These colonies were collected from twigs on the forest floor. During intensive field collection, five mature colonies containing female adultoids with elongated abdomens, that is, mature adultoid queens, were successfully collected (Table S1). Mature adult queens were also included in the ploidy analysis. Ploidy analysis was performed on 15 female alates, 15 young adultoid queens, three field-collected young adultoid queens, and five mature field-collected adultoid queens.

**Queen fecundity and fresh body weight of queens**

To examine the relationship between fresh body weight and fecundity of termite queens, we obtained nymphoid queens of various body weights from six colonies (A–F) collected from pine or cedar forests in Kyoto, Shiga, and Wakayama from June to July of 2016 and 2017. Five mature nymphoid queens from A, D, E, and F; six from B, ten from C were chosen. Their fresh body weights were measured with a digital balance in milligrams to two decimals. Thirty-six queens were isolated from their nests and kept in the darkness for five days with 100 workers from the same nest. As an experimental nest, we used a 30 mm Petri dish lined with moist unwoven cloth, which was their food material [55]. After five days, the number of oviposited eggs was counted. To evaluate the validity of the fresh body weight of the collected queens as a proxy for fecundity, we used a GLMM with Poisson errors. In this model, the number of eggs oviposited for five days was the response variable, and the initial fresh body weight was the explanatory variable. The original colony was included as the random effect.

**Termite samples for the image-based ploidy analysis on the fat body**

To obtain workers, nymphs, and laboratory-emerged nymphoid queens, we collected three field colonies (HM210907A, HM210907B, and OZ210902) without royals from pine forests in Aichi and Shizuoka, on September 2021. Workers, 4th instar nymphs, and young nymphoid queens with undeveloped ovaries were carefully extracted and used immediately in the experiment. These young nymphoids likely emerged after the nest wood was transported to the laboratory, potentially because they were orphaned at the time of collection. The remaining termites comprising workers and nymphs were used to establish large-scale orphaned colonies (Fig. S3). These cases were maintained at 25 °C in constant darkness for three months. The cases were carefully opened, and individuals were extracted. Egg piles were observed in all three established orphan colonies, indicating that the emerged nymphoid queens were reproducing.

**Cell sorting and microscopic observation for the image-based ploidy analysis on the fat body**

The fat bodies and heads of five reproducing nymphoid queens from orphaned colonies derived from HM210907A and HM210907B were dissected in PBS and processed for ploidy analysis using the method described in the main text (see “Ploidy analysis of the fat body of termites using flow cytometry” in Methods). During dissection, the ovaries of these queens were confirmed to contain vitellogenic oocytes. Stained nuclei were analyzed for DNA-PI fluorescence using a Cell Sorter SH800 (SONY Imaging Products and Solutions) with a Sorting Chip (100 μm) at an excitation wavelength of 488 nm. Based on the signal intensity of the DNA-PI fluorescence, we identified and gated populations with 2C, 4C, and 8C using the Cell Sorter Software (ver. 2.1.5). We sorted the nuclei with each ploidy level into 1.5 mL Eppendorf tubes, using the “normal sorting” mode. The sorted nuclei were stained again with PI (1 μg/mL, Dojindo) and visualized using fluorescence microscopy. Nuclei were imaged using a BX-61 microscope (Olympus) and a DS-Fi1 CCD camera (Nikon). Images were processed using ImageJ software (National Institutes of Health, USA, http://rsb.info.nih.gov/ij/). The diameter of nuclei with each ploidy level were measured using the “Particle analysis” function of the software. We evaluated the validity of the size of nuclei in queens as a proxy for ploidy levels. The sizes of nuclei that were sorted as 2C nuclei from head samples and 2C, 4C, and 8C nuclei from fat body samples were compared using LMM followed by Tukey’s HSD post hoc test. In this analysis, ploidy levels and body parts were fixed effects and the original colony was included as a random effect.

**SI table and figures**

**Table S1.** Sample information of *R. speratus* colonies used for ploidy analysis

| Colony code* | Sampling location | Usage (# of individuals used)^†^ |
| --- | --- | --- |
| YS180815A | Yamashina-ku, Kyoto, Kyoto | FCM [female nymph (3), nymphoid queen 0 day (3)] |
| YS180815B | Yamashina-ku, Kyoto, Kyoto | FCM [female nymph (3), nymphoid queen 0 day (3)] |
| YO180916 | Sakyo-ku, Kyoto, Kyoto | FCM [female nymph (3), nymphoid queen 0 day (3)] |
| HI150720 | Hiei-daira, Ohtsu, Shiga | FCM [nymphoid queen 3 month (5), ergatoid queen 3 month (5)] |
| UR150715A | Sakyo-ku, Kyoto, Kyoto | FCM [nymphoid queen 3 month (5)] |
| YO150715B | Sakyo-ku, Kyoto, Kyoto | FCM [nymphoid queen 3 month (5)] |
| YO151006 | Sakyo-ku, Kyoto, Kyoto | FCM [nymphoid queen 0 day (3), ergatoid queen 0 day (6)] |
| UR151008 | Sakyo-ku, Kyoto, Kyoto | FCM [nymphoid queen 0 day (3), ergatoid queen 0 day (4)] |
| TA151014 | Sakyo-ku, Kyoto, Kyoto | FCM [nymphoid queen 0 day (3)] |
| GB150703 | Enjugahama, Gobo, Wakayama | FCM [ergatoid queen 3 month (5)] |
| OK150625B | Sakyo-ku, Kyoto, Kyoto | FCM [ergatoid queen 3 month (5)] |
| IN150509B | Inagawa, Kawabe, Hyogo | FCM [female alate (5), colony founding adultoid queen (3)] |
| FU150426B | Inagawa, Kawabe, Hyogo | FCM [female alate (5), colony founding adultoid queen (3)] |
| RU150510Z | Sonobe, Nantan, Kyoto | FCM [female alate (5), colony founding adultoid queen (3)] |
| OO150510 | Sakyo-ku, Kyoto, Kyoto | FCM [field-collected mature nymphoid queen (8)] |
| OK150625 | Sakyo-ku, Kyoto, Kyoto | FCM [field-collected mature nymphoid queen (10)] |
| OK150630 | Sakyo-ku, Kyoto, Kyoto | FCM [field-collected mature nymphoid queen (10)] |
| SA150704A | Tamba-sasayama, Hyogo | FCM [field-collected mature nymphoid queen (10)] |
| SA150704B | Tamba-sasayama, Hyogo | FCM [field-collected mature nymphoid queen (10)] |
| SA150704C | Tamba-sasayama, Hyogo | FCM [field-collected mature nymphoid queen (10)] |
| SA150704D | Tamba-sasayama, Hyogo | FCM [field-collected mature nymphoid queen (9)] |
| OK150715 | Sakyo-ku, Kyoto, Kyoto | FCM [field-collected mature nymphoid queen (10)] |
| MI150719E | Sakyo-ku, Kyoto, Kyoto | FCM [field-collected mature nymphoid queen (10)] |
| NI150719 | Chuoku, Niigata, Niigata | FCM [field-collected mature nymphoid queen (10), young nymphoid queen (4)] |
| PO150806H | Nishikyo-ku, Kyoto, Kyoto | FCM [field-collected mature nymphoid queen (8)] |
| OH150806C | Nishikyo-ku, Kyoto, Kyoto | FCM [field-collected mature nymphoid queen (10)] |
| HI160412 | Sakyo-ku, Kyoto, Kyoto | FCM [field-collected young nymphoid queen (5)] |
| TJ150418B | Tanakamisato, Otsu, Shiga | FCM [field-collected mature adultoid queen (1)] |
| GB150703B | Mihama, Hidaka, Wakayama | FCM [field-collected mature adultoid queen (1)] |
| OH150806A | Nishikyo-ku, Kyoto, Kyoto | FCM [field-collected mature adultoid queen (1)] |
| IW160518 | Sakyo-ku, Kyoto, Kyoto | FCM [field-collected mature adultoid queen (1), mature nymphoid queen (3), young nymphoid queen (1)] |
| MK160809 | Matsushiro, Nagano, Nagano | FCM [field-collected mature adultoid queen (1)] |
| HI150720A | Sakyo-ku, Kyoto, Kyoto | FCM [field-collected founding adultoid queen (1)] |
| HI150720B | Sakyo-ku, Kyoto, Kyoto | FCM [field-collected founding adultoid queen (1)] |
| HI150720C | Sakyo-ku, Kyoto, Kyoto | FCM [field-collected founding adultoid queen (1)] |
| HM210907A | Minami-ku, Hamamatsu, Shizuoka | CS [nymphoid queen 3 month (5)], CA [female worker (1), female nymph (1), nymphoid queen 0 day (1), nymphoid queen 3 month (1)] |
| HM210907B | Minami-ku, Hamamatsu, Shizuoka | CS [nymphoid queen 3 month (5)] |
| OZ210902 | Okuyamada, Okazaki, Aichi | CA [female worker (1), female nymph (1), nymphoid queen 0 day (1), nymphoid queen 3 month (1)] |

^*^Numbers in “colony code” indicate the dates when the colonies were collected. For example, colony YS180815A was collected on April 5, 2018.

^†^Abbreviations: FCM, ploidy analysis using flow cytometry; CS, cell sorting and nuclear size analysis; CA, cell type and ploidy analysis using an image-based method.

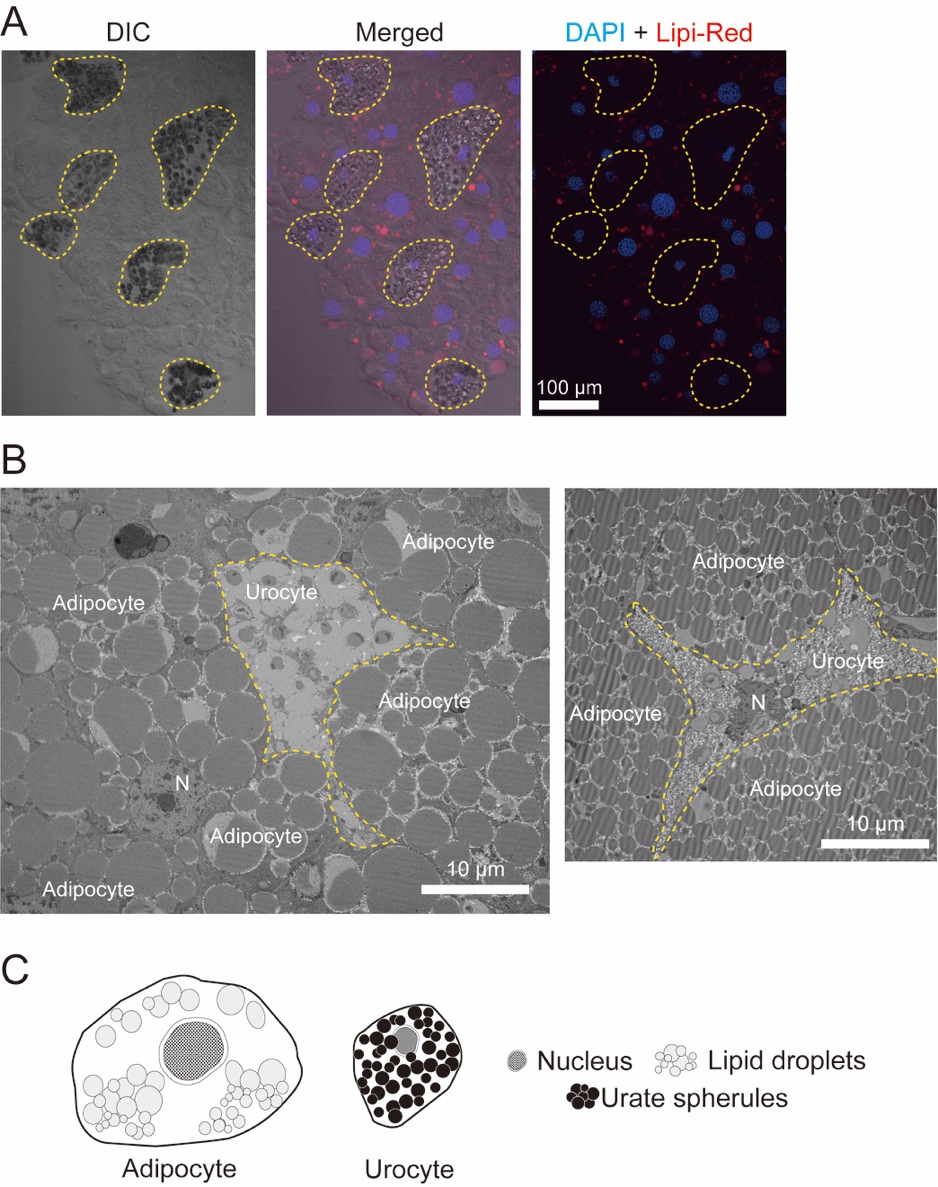

**Figure S1.** Cell types in the termite fat body. **A** Morphology of fat body cells of a worker under differential interference contrast (DIC) and confocal microscopy. DNA and lipid dropet were stained using DAPI (blue) and Lipi-Red (red), respectively. In the DIC images, urate spherules appeared to be reflective particles. Nuclei of cells that contained lipid droplets (adipocytes) were more circular than those surrounded by urate spherules, and this was urocyte. Urocytes were enclosed by yellow dashed lines. **B** Transmission electolon microscopy for the fat body of a field collected queen (left) and worker (right). Cells that had urate spherules with spherical concretions that appeared to be empty or with a small dense central core and that contained lipid droplets were considered as urocytes and adipocytes, respectively (according to [23]). The urocyte was enclosed by yellow dashed lines. N, nucleus. **C** Illustration of the two types of cells in the termite fat body, which was drawn based on the observations from this study.

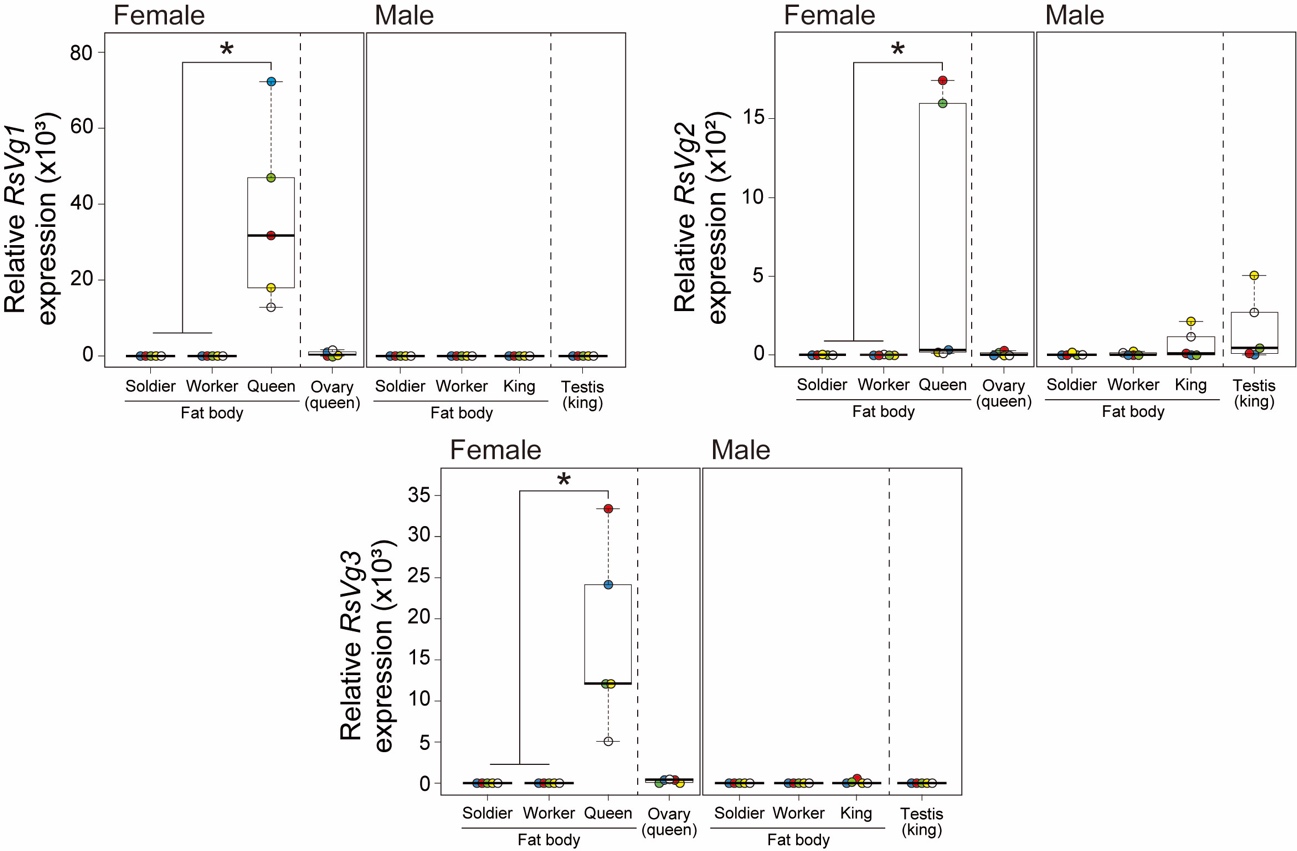

**Figure S2.** Vitellogenin gene expression in the termite queen fat body. The expression levels of vitellogenin genes (*RsVg1*–*3*) were measured using qPCR in the fat body of both sex of workers, soldiers and reproductives, and reproductive organs, that is, the ovary of the queen and testis of the king. Relative expression levels were calculated using a typical ΔΔCt method relative to the fat body of female workers. Box plots show the median (center line), 75th percentiles (top of box), 25th percentiles (bottom of box), and whiskers connecting the largest and smallest values within 1.5 interquartile ranges. Each color is corresponding to the derived colony (n = 5). The queen fat body exhibited significantly higher expression levels of all the three genes than those of female workers and soldiers (Mann–Whitney U tests, Bonferroni correction, **p* < 0.05).

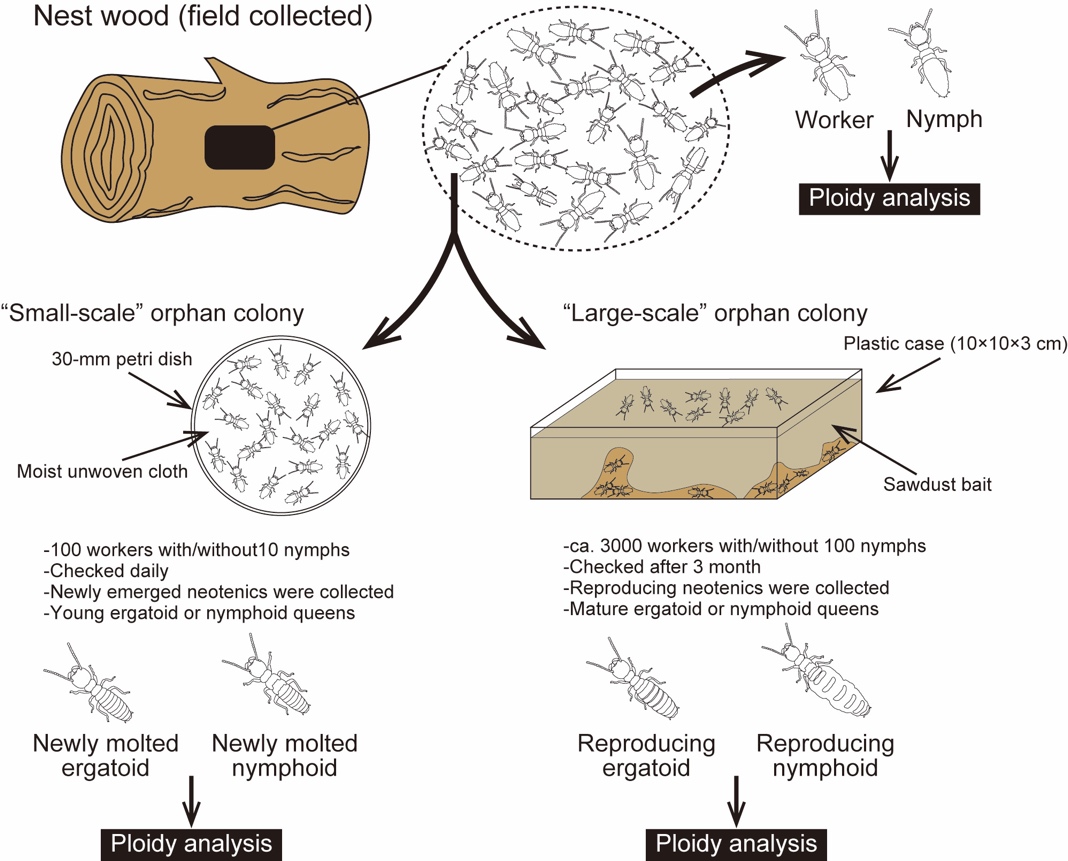

**Figure S3**. Establishment of artificial colonies under laboratory conditions. The nest woods collected from the field were carefully destroyed, and individuals including workers, soldiers and nymphs were extracted. Workers and 4^th^ instar nymphs were processed on the day and the ploidy level in their fat body was analyzed. Other workers and nymphs were assigned to two types of orphan colonies. In the “small-scale” orphan colony, 100 workers or 100 workers with 10 nymphs were placed on a 30-mm Petri dish lined with moist unwoven cloth as food and were checked daily. When workers and nymphs differentiated into neotenics, that is, ergatoids and nymphoids, respectively, the individuals were sexed, and females were processed on the day for ploidy analysis. For the “large-scale” orphan colony, approximately 3000 workers or 3000 workers with 100 nymphs were placed on a plastic case filled with sawdust baits to obtain reproducing neotenics. After three months, the termites were carefully extracted, and the queens were processed for ploidy analysis.

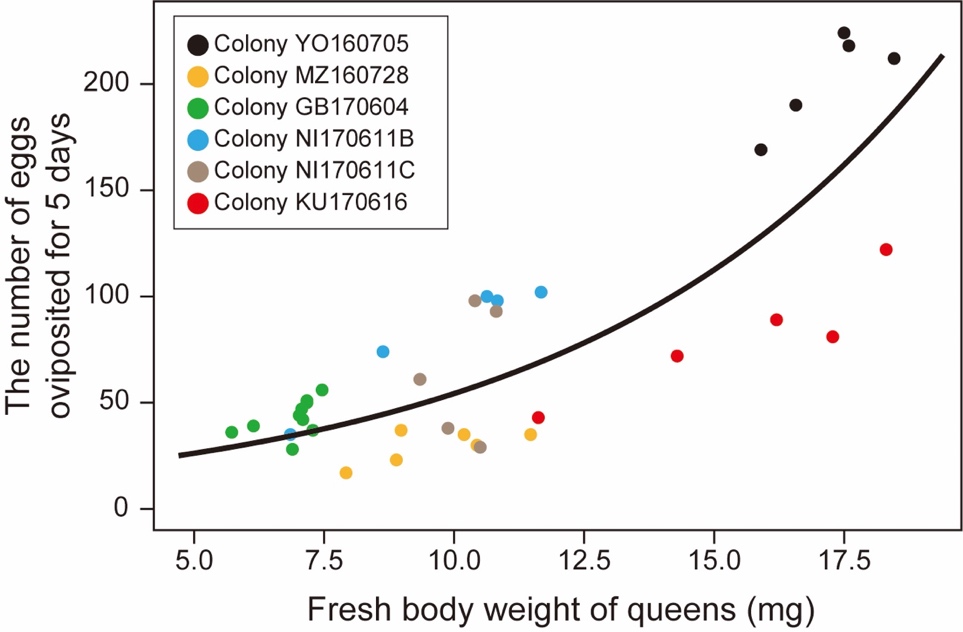

**Figure S4.** Effects of initial body weight of queens on fecundity for five days. Dots are individual values, and their colors indicate original colonies. Analysis using a generalized linear mixed model (GLMM) was conducted for the dataset of egg production for five days and initial weight of queens from five colonies. The weight of queens had significant effect on the queen fecundity, that is, the number of eggs produced for five days (GLMM with type II Wald chi-square test, *χ*^2^ = 97.308, *df* = 1, *p* < 0.01). The black line is the regression curve calculated with GLMM, and the formula:

$$y = e^{(0.1457x + 2.53750)}$$

The correlation value of the fixed effect, the initial body weight was −0.753.

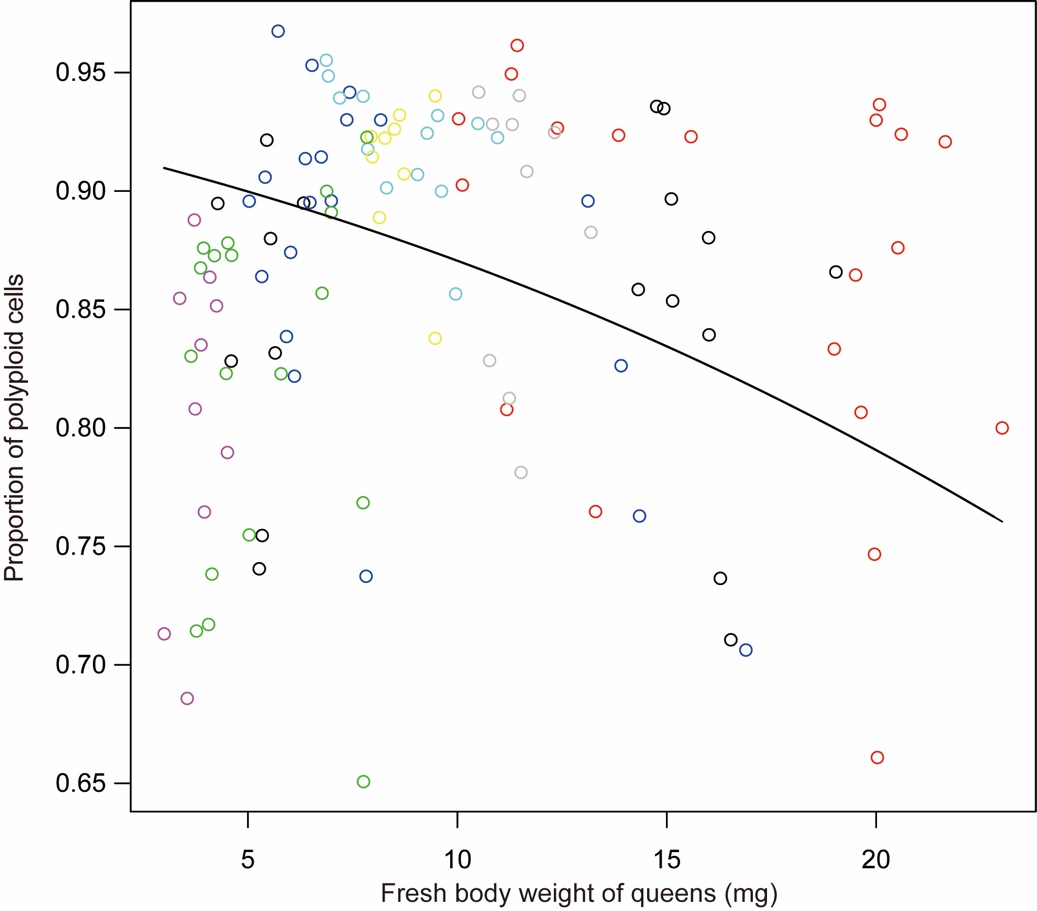

**Figure S5.** Relationship between fresh body weight of queens and the proportion of polyploid cells in the fat body. Dots are individual values, and their colors indicate original colonies. Analysis using a generalized linear mixed model (GLMM) was conducted for the dataset of field collected mature queens from May to August. The weight of queens had a significant effect on the proportion of polyploid cells in the fat body (GLMM with type II Wald chi-square test, fresh body weight: *χ*^2^ = 109.71, *df* = 1, *p* < 0.01). The black line is the regression curve calculated with GLMM, and the formula:

$$y = \frac{1}{{1+e}^{-(-0.1651 x + 2.7792)}}$$

The correlation value of the fixed effect, the fresh body weight was relatively low, −0.360.

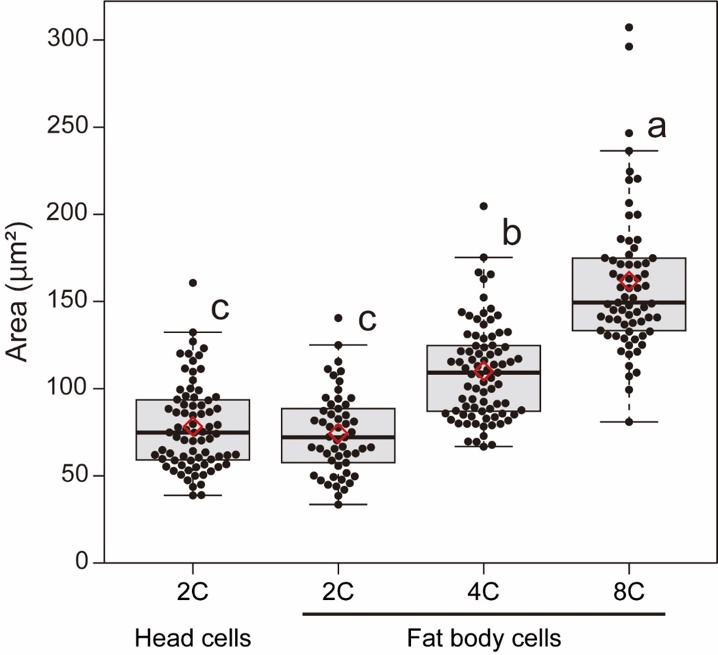

**Figure S6** Size distribution of sorted nuclei from the head and fat body of nymphoid queens. Box plots show the median (center line), 75th percentiles (top of box), 25th percentiles (bottom of box), and whiskers connect the largest and smallest values within 1.5 interquartile ranges. The red diamonds indicate mean percentages of each category. Dots are individual values. Different letters indicate significant differences among means in each comparison group (Tukey’s HSD test, *p* < 0.05).

**References (only referred to in the SI)**

51. Tasaki, E., Komagata, Y., Inagaki, T., & Matsuura, K. (2020). Reproduction deep inside wood: a low O2 and high CO2 environment promotes egg production by termite queens. Biology Letters, 16(4), 20200049.
52. Weesner, F. M. (1969). The reproductive system. In K. Krishna, & F. M. Weesner (Eds.), Biology of termites (vol. I, pp. 125–160). New York, NY: Academic Press.
53. Zimet, M., & Stuart, A. M. (1982). Sexual dimorphism in the immature stage of the termite, *Reticulitermes flavipes* (Isopreta: Rhinotermitidae). Sociobiology, 7, 1–7.
54. Nozaki, T., Yashiro, T., & Matsuura, K. (2018). Preadaptation for asexual queen succession: queen tychoparthenogenesis produces neotenic queens in the termite *Reticulitermes okinawanus*. Insectes Sociaux, 65(2), 225–231.
55. Nozaki, T., & Matsuura, K. (2021). Oocyte resorption in termite queens: Seasonal dynamics and controlling factors. Journal of Insect Physiology, 131, 104242.
